# Identifying differentially methylated sites in samples with varying tumor purity

**DOI:** 10.1101/248781

**Authors:** Antti Häkkinen, Amjad Alkodsi, Chiara Facciotto, Kaiyang Zhang, Katja Kaipio, Sirpa Leppä, Olli Carpén, Seija Grénman, Johanna Hynninen, Sakari Hietanen, Rainer Lehtonen, Sampsa Hautaniemi

## Abstract

DNA methylation aberrations are common in many cancer types. A major challenge hindering comparison of patient-derived samples is that they comprise of heterogeneous collection of cancer and microenvironment cells. We present a computational method that allows comparing cancer methylomes in two or more heterogeneous tumor samples featuring differing, unknown fraction of cancer cells. The method is unique in that it allows comparison also in the absence of normal cell control samples and without prior tumor purity estimates, as these are often unavailable or unreliable in clinical samples. We use simulations and next-generation methylome, RNA, and whole-genome sequencing data from two cancer types to demonstrate that the method is accurate and outperforms alternatives. The results show that our method adapts well to various cancer types and to a wide range of tumor content, and works robustly without a control or with controls derived from various sources. The method is freely available at https://bitbucket.org/anthakki/dmml.

## 1 Introduction

Aberrant DNA methylation is a hallmark of all cancer types (Hanahan and Weinberg, 2011; Witte *et al*., 2014; Shen and Laird, 2013; Timp and Feinberg, 2013). Compared to genomic alterations, DNA methylation offers a more flexible yet a persistent mechanism to exert changes on the phenotype, which manifests in silencing tumor suppressor genes, activating proto-oncogenes, or causing chromosomal instability (Witte *et al*., 2014; Shen and Laird, 2013; Timp and Feinberg, 2013; Yang *et al*., 2015). While some patterns have been identified, the role of DNA methylation alterations in cancer development, tumor pathogenesis, and treatment response varies between cancer types (Witte *et al*., 2014; Yang *et al*., 2015; Ciriello *et al*., 2013). Advances in nextgeneration sequencing (NGS) in combination with classical bisulfite conversion (Frommer *et al*., 1992; Harris *et al*., 2010) have allowed profiling methylomes at a single nucleotide resolution at an unprecedented scale (Shen and Laird, 2013; Ciriello *et al*., 2013). These developments have surged an interest to develop personalized clinical applications that employ DNA methylation alterations as diagnostic and prognostic biomarkers and as therapeutic targets (Wei *et al*., 2006; Altman *et al*., 2013). A major challenge in this is that the surgically removed samples comprise of heterogeneous mixture of cancer cells and the microenvironment. As the exact tumor content (tumor cell fraction, tumor purity) and cell composition varies considerably between the samples, direct comparison of samples even from the same patient without correcting for the tumor cell content can lead to spurious results (Carter *et al*., 2012; Aran *et al*., 2015; Zheng *et al*., 2017).

Immunohistochemical staining has been the most used technique to determine cell composition in tissue sections and to select high purity samples for further experiments. More recently, high-throughput measurement technologies have enable cost-efficient and rapid production of patient-derived molecular data, but also allow estimating the sample tumor content. Computational tools have been developed for purity estimation using somatic copy-number data (Carter *et al*., 2012; Oesper *et al*., 2013; Van Loo *et al*., 2010), single-nucleotide polymorphisms (Van Loo *et al*., 2010), variant allele frequency of somatic mutations (Carter *et al*., 2012), or RNA expression (Yoshihara *et al*., 2013). Evidence suggests that these methods outperform manual analysis and allow analysis of large-scale datasets (Carter *et al*., 2012; Zheng *et al*., 2017). While the tools allow fast and reproducible tumor purity analysis, most differential DNA methylation analysis methods i) do not account for the sample heterogeneity or adjust the analysis for low tumor purity or differences between samples (Hebestreit *et al*., 2013; Hansen *et al*., 2012; Feng *et al*., 2014; Sun *et al*., 2014; Sun and Yu, 2016; Wang *et al*., 2016), ii) require a library of control samples (Houseman *et al*., 2012), or iii) do not account for co-methylation of closely located sites, cannot correct for this at singlenucleotide level (Hebestreit *et al*., 2013; Hansen *et al*., 2012; Feng *et al*., 2014), or model co-methylation uniformly (Sun and Yu, 2016; Wang *et al*., 2016; Zheng *et al*., 2014). All the shortcomings lead to biased findings and false biological interpretation. The requirement for a controls is problematic, as in many cases appropriate controls are not available or are incomparable, and the purities predicted using matched blood controls show poor correlation with the estimates derived with universal normal controls in various cancer types (Zheng *et al*., 2017).

Here, we present a method based on a latent stochastic model which allows comparing DNA methylomes at a single-nucleotide resolution between two or more tumor samples with different, unknown tumor purities. The method performs accurately without a normal cell control sample or prior tumor purity estimates, and accounts for spatially comethylated cytosines, improving accuracy at lower coverage sites. We demonstrate the superior performance using simulations and two sets of NGS cancer data — targeted sequencing data from high-grade serous ovarian cancer patients and genome-wide reduced representation bisulfite sequencing data from diffuse large B-cell lymphoma patients.

## 2 Models and methods

Patient-derived samples are composed of different cell types, such as various types of stromal and immune cells in addition to cancer cells, that are present in different, unknown proportions. The number of cell types that can be identified is limited by the setting, e.g. a pairwise comparison of two tumors can be corrected for a single common nuisance. We use sequencing data to estimate the underlying DNA methylome of each cell type and the composition of each sample simultaneously. These estimates allow testing cancer cell specific differences between the samples and obtaining purity-corrected estimates of the methylomes of each cell type. An overview is shown in Figure 1, and full details are given in Supplementary material.

**Figure 1:**
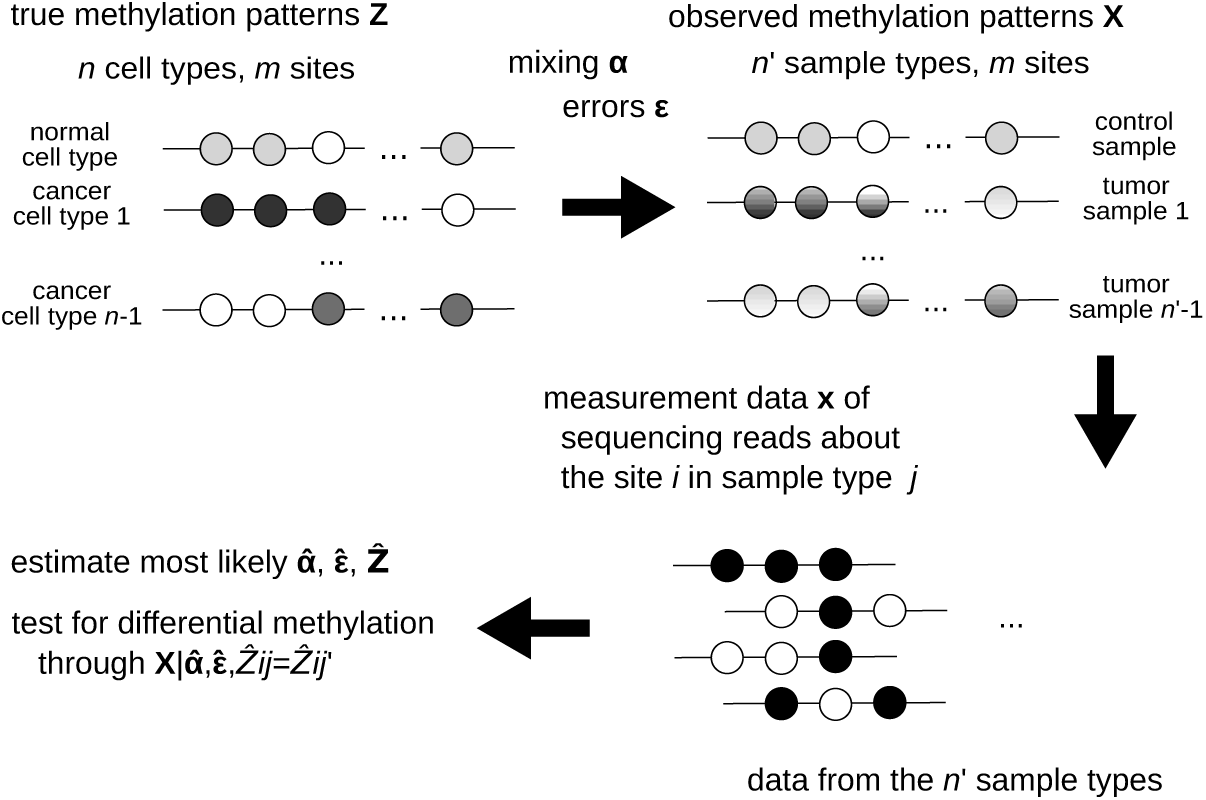
Overview of the proposed methodology. The objective is to identify features of the true methylation patterns of *n* cell types at *m* sites. We assume a generative model where each of the *n*′ sample types is a mixture of the *n* cell types, with unknown proportions and unmodeled additional variations (errors). Sequencing reads from each of the *n*′ sample types are pooled and grouped into w site runs for local intersite modeling. These data are used to estimate the most likely model, which can be used to test if a pair of cell types feature differential methylation.

### 2.1 Modeling latent methylation patterns

We assume that there are *n* (pure) cell types, *j*:th of which has an unknown methylation pattern 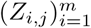 defined at *m* sites. In a typical case comparing two tumor samples, three hypothesized cell types exist: a single normal cell type, common in both samples, and two cancer cell types, (potentially) unique in each tumor sample (cf. Figure 1).

The *m* sites at which the methylation is modeled can represent either adjacent or distant genomic sites, making the method applicable to whole-genome bisulfite sequencing (WGBS), targeted bisulfite sequencing (TBS), reduced representation bisulfite sequencing (RRBS) (Harris *et al*., 2010; Lee *et al*., 2013), or other sequencing-based platforms. For methylation, the latent state variables *Z*_*i*,*j*_ are binary, but could be e.g. quaternary for detecting DNA mutations.

### 2.2 Modeling observed methylation patterns

A typical tumor sample consists of few major cell types, such as stromal and immune cells in addition to the actual cancer cells of interest (Carter *et al*., 2012; Aran *et al*., 2015). Due to this, there is a distinction between the methylation patterns that are measured in the impure samples and those of the underlying pure cell types (which are not directly measured).

Each sample is assumed to be a mixture of the *n* cell types and be further corrupted by random noise (cf. Figure 1). We use *X*_*i*,*j*_ to denote the random variable corresponding to the methylation read at site *i* of the *j*:th sample type, *α*_*k*,*j*_:s to denote the mixing parameters and *ϵ*_*k*,*j*_ to denote the per-site error rate (cf. Figure 1). The mixing parameter *α*_*k*,*j*_ determines the fraction of cells of type *k* in the sample *j*, which will be determined by the unknown sample composition. The error rates depend on various factors such as unmodeled variations for the cell type *k* and sequencing and postprocessing errors of the sample *j*, which makes their direct measurement cumbersome, but this is not a problem as the parameters can be estimated from the data. Here, the errors are considered to be independent random bit-flips, but a more complex error model is possible (see Supplementary material).

In the simple case of comparing two tumor samples, we use a single common error parameter *ϵ*_*k*,*j*_ = *ϵ*, three cell types (*n* = 3) as described in the previous section, and three sample types (*n′* = 3): a normal cell sample (control) and two kinds of tumor samples. The mixing is determined by the two mixing parameters *α*_1,1_ = *α*_1_ and *α*_1,2_ = *α*_2_, the tumor purities of each two tumor samples, and the other mixing parameters are implicit: *α*_0,0_ = 1, *α*_*kl*,*1*_ = 0 (control is pure), *α*_*0*,*j*_ = 1 – *α*_*j*,*j*_ (the impurities in the tumor samples are normal cells) and *α*_2‒*j*+1,*j*_ = 0 (no crosstalk between the two cancer cell types).

An arbitrary number of cell types and sample types are supported, provided that the problem is identifiable (e.g. it is not possible to decompose a single sample into multiple cell types without further constraints). For example, an arbitrary number of tumor samples can be compared simultaneously, provided that the normal cells feature a similar methylation pattern in each tumor sample. Alternatively, multiple normal cells can be present, provided that an appropriate number of controls are supplied. The identifiability problems arising from low sample size are locally mitigated by the co-methylation modeling (see next section).

### 2.3 Co-methylation modeling

In general, even for RRBS, a single sequencing read will contain multiple adjacent sites (in CpG islands, we expect about 3 to 4 CpG sites per a 100 basepair read (Illingworth and Bird, 2009)), which introduces correlations in the mixing. This, and the fact that alterations in methylation patterns (Eckhardt *et al*., 2006) tend to span larger regions, introduces correlations in the adjacent sites *X*_*i*,*j*_ and *X*_*i*′,*j*_, which result in incorrect statistical significance and wrong calls. Note that while the latter type of correlations are amplified in RRBS and using WGBS instead would dilute such correlations, the former type of correlations would be amplified.

To capture these correlations, we model *w*-wise joint distributions of the methylation patterns. We found this approach to be the most appropriate, as sufficiently distant sites are expected to be uncorrelated, while the correlation between adjacent sites varies e.g. depending on distance (for RRBS and targeted sequencing) and boundaries of genetic elements (Eckhardt *et al*., 2006; Lister *et al*., 2009), calling for local modeling in each neighborhood. Using a larger window size *w* increases computational effort but has no other disadvantages: if the data lacks dependence, the results are equal to those using a smaller window size.

### 2.4 Model estimation and differential methylation testing

Given the above model, our objective is to use the sequencing data to estimate the underlying methylation pattern of each (pure) cell type and the composition of each tumor sample type simultaneously.

The estimation is performed in maximum likelihood sense (ML; i.e. select parameters that most likely generate the data), which is implemented through a numerical expectation maximization algorithm (see Supplementary material). Direct optimization is infeasible even for moderate number of sites as the complexity is 𝒪(|∑|^*mn*^) where *m* and *n* are the number of sites and cell types, respectively, and |Σ| = 2 is the alphabet size. Meanwhile, our method is exponential time and space in *w n*, where *w* is the window size, but takes linear time (per iteration) and constant memory in *m*, which allows genome-scale analysis. Prior information about the parameters can be included by modifying the ML objective. For example, if estimates of the tumor purities are available from other sources, the estimator can be persuaded to use this information.

The estimates allow testing if specifically the cancer cells in the different sample types feature a differential methylation: we derive a p-value through a likelihood ratio test (see Supplementary material). Alternatively, the estimated distribution of the cell type methylation patterns can be used to derive maximum posterior estimates of the cell type methylation patterns, yielding “purified” methylation counts.

## 3 Results and discussion

### 3.1 Monte Carlo simulations

The use of sequencing data from cancer patients allow estimation of sensitivity to some degree but not specificity. Therefore, a simulation where ground truth is known is important. Here, we employed Monte Carlo simulations based on publicly available WGBS data and compared differential DNA methylation calling under varying tumor purity.

#### 3.1.1 Simulation settings

We obtained sequences of a length *m*, for three cell types (one normal and two cancer cells; the simplest setting where multiple tumor samples are compared) by sampling publicly available whole-genome bisulfite sequencing (WGBS) data of immortalized cell lines from The ENCODE Project Consortium (2012) (see Supplementary material). Afterwards, bisulfite reads were generated by sampling the reads of the corresponding WGBS dataset, extracting the methylation signal, and adding random errors (independent bit flips). The control sample was generated from the GM12878 cell line, while two tumor samples were generated by mixing the K562 and HepG2 cancer cell lines with the GM12878 line.

Methylation calls were made at a significance level 0.05 after adjusting for false discovery rate (FDR) using the Benjamini-Hochberg procedure (Benjamini and Hochberg, 1995), following the procedure of Lister *et al.* (2009). An FDR adjustment and the selected significance level strongly influence the number of true versus false calls, but we did not find it to affect our conclusions (not shown).

#### 3.1.2 Performance under varying tumor purity

First, we studied how varying sample purities affects the performance of detecting differentially methylated sites in two tumor samples. We compared our approach with several existing methods: Fisher’s exact test (Lister *et al*., 2009; Pan *et al*., 2015; Assenov *et al*., 2016), MOABS (Sun *et al*., 2014), a mixture-adjusted Fisher’s exact test (see Supplementary material), InfiniumPurify (Zheng *et al*., 2017), and DSS (Feng *et al*., 2014). In the comparisons, we used our method with and without a control sample and using various window sizes *w* for the co-methylation modeling. Unlike our method, none of the alternatives can perform control-free differential methylation calling on two tumor samples without known purities, which is a severe limitation, and in the comparisons, they are used with additional information. Most of the alternative methods for detecting differential methylation are unsuitable for a setting comparing two or more heterogeneous tumor samples, as they assume that the samples are pure or feature similar purity (Hebestreit *et al*., 2013; Hansen *et al*., 2012; Sun and Yu, 2016; Wang *et al*., 2016), or cannot compare multiple tumor samples (e.g. but only tumor versus normal) (Zheng *et al*., 2014).

Figure S1 exemplifies a typical setting with 30× average coverage with *m* = 1,000 CpG sites (chromosome 6, 22,333,542-22,469,062 in GRCh38) and sample purities of *α*_*1*_ = 0.25 and *α*_*2*_ = 0.75 and an error rate of 0.05. As expected, Fisher’s exact test performs poorly — not much better than a random guess — due to the different sample purities, while the mixture-adjusted variant can reach an accuracy of about 90%. MOABS performs comparably to the Fisher’s exact test, and InfiniumPurify and DSS comparably to the mixture-adjusted Fisher’s exact test, which is expected as the latter set of methods properly model the tumor composition while MOABS does not. Meanwhile, our method provides higher accuracy than the alternatives over a wide range of thresholds, and generally ranks higher in either (or both) specificity or sensitivity. Using a matched control and higher order co-methylation modeling results in further improvements in the performance, which are unavailable to the alternatives.

For a more thorough analysis, we simulated data with varying sample purities in each sample. The results were obtained with 30× average coverage, and the average false call rates of 500 simulations with *m* = 1,000 CpG sites each as a function of the sample purities are visualized in Figure 2. The lower triangle of each heatmap shows the fraction of false negatives, and the upper triangle the total fraction of false calls (false negatives plus false positives; the fraction of false positives ought to be small as they are being controlled for). All methods perform poorly (equal to random) at purities *a*_1_ = 0 or *a*_2_ = 0. This is expected, as the samples do not contain any information about the methylation patterns of the cancer cells in one (or both) of the samples. Compared to the alternatives, our methods develop these false calls at a later stage. Meanwhile, when the purities are dissimilar, Fisher’s exact test and MOABS generate a large amount of false positives, which is the major reason for their unsuitability for these data. Our method, when used without a control, is susceptible to generating false negatives when one of the samples is pure and the other is not (i.e. *a*_1_ ~ 1, *a*_2_ ~ 0.5 or *a*_1_ ~ 0.5, *a*_2_ ~ 1), which is due to the fact that it is not possible to identify which of the patterns in the impure sample are from normal and which are from cancer cells, as the pure sample lacks the normal cells. When a control is provided, there is no such ambiguity, which is also why this issue does not appear with the other methods. Results with higher coverage qualitatively similar, but the accurate range of purities (e.g. 95% accuracy) is increased for each method.

**Figure 2:**
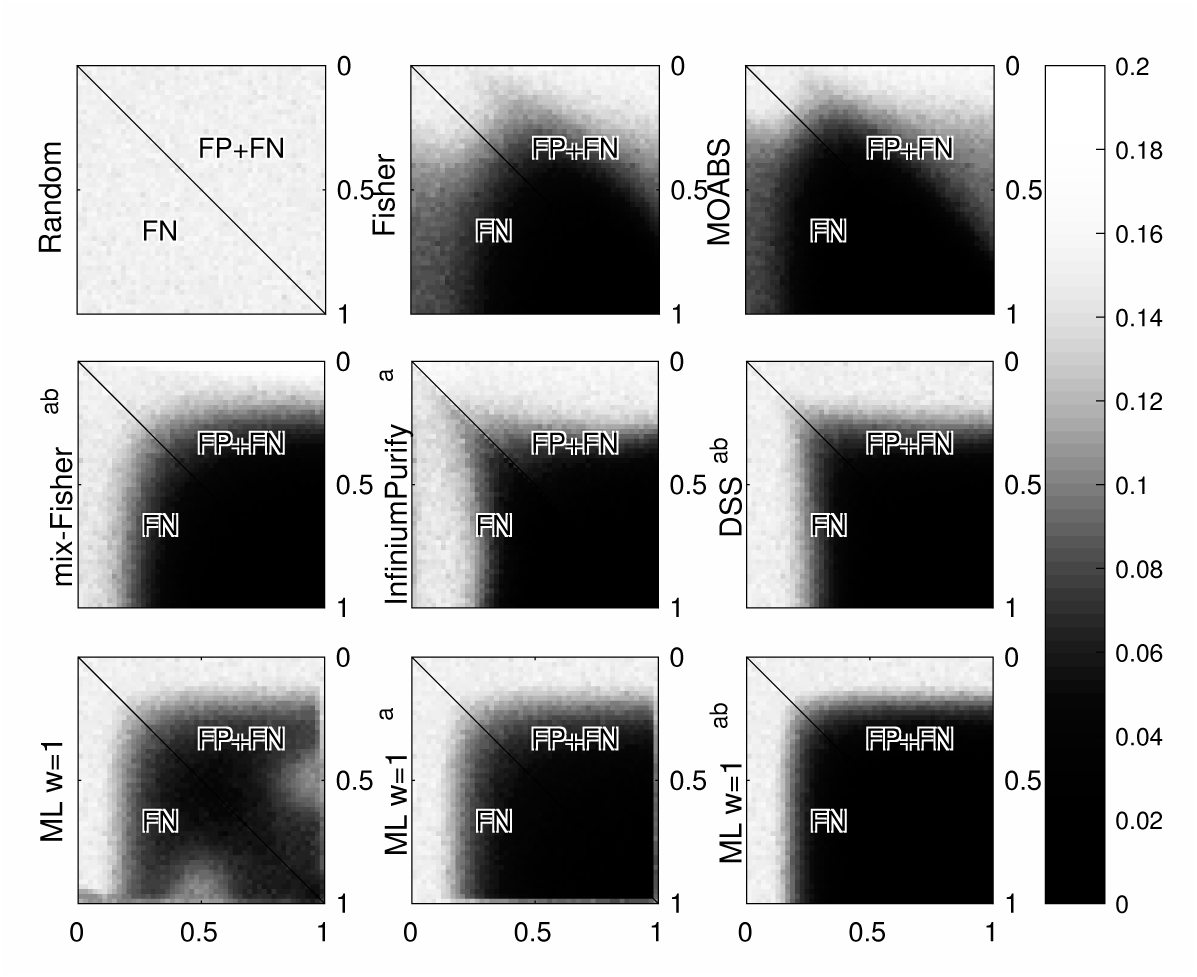
False negatives/false calls with simulated data with varying sample purities. The axes represent tumor purity of the two samples to be compared and the heatmap intensity represents the average false negative rate (FN; lower triangle) and total false calls rate (FN+FP; upper triangle) in 500 simulations. Different panels represent different methods: Fisher’s exact test (Fisher), MOABS, mixture-adjusted Fisher's exact test (mix-Fisher), InfiniumPurify, DSS, our maximum likelihood method (ML) with a window size *w* = 1. Methods with ^a^ use a matched control sample, and methods with ^b^ the true purities.

In Figure S2, we show how the performance varies across the simulations. This Figure Shows the distribution of false calls in the 500 simulations for each method along the curve *α*_1_ = 1 — *α*_2_. The results indicate that our methods perform competitively for various degrees of tumor purity, and that providing the control or prior information about the purities offers further advantages. In general, providing a prior results in more consistent performance (lower variance), while providing the control mainly enhances the average performance.

#### 3.1.3 Advantages of co-methylation modeling

Next, we show that our method offers even better performance, provided that the sequencing reads are of sufficient length. This feature allows our method to have good performance even in low coverage settings provided that the reads span over multiple CpG sites, which is advantageous especially with targeted sequencing data. The alternatives do not exploit this information. Similarly, our method with a window size of *w* = 1 lacks correlation modeling, rendering this information unexploited.

An example with 30× average coverage in a 1,000 site experiment, with *α* = 0.25, *α*_2_ = 0.75, and an error rate of 0.05, collected from 500 simulations is summarized in Figure S3. For a read length of exactly 1 CpG site, the results with *w* > 1 are equal to that with *w* = 1. However, when a single read covers multiple CpG sites, the co-methylation modeling allows for a greater accuracy. Further increases in the window size offer additional improvements in the accuracy. The data that were used as a basis of the simulation features about 4 CpG sites per read on average, so we are unable to show the advantages beyond this read length, but fully synthetic simulations (not shown) suggest that further improvements are possible for larger read lengths as well.

### 3.2 Ovarian cancer dataset

Next, we applied the various methods to detect differential methylation between samples surgically removed from patients diagnosed with high-grade serous ovarian cancer (HGSOC). HGSOC is responsible for more than 40,000 deaths annually in Europe alone and more than 50% of the patients die within five years of diagnosis (Berns and Bowtell, 2012).

#### 3.2.1 Sample description

We used a total of five ovarian cancer tumor samples from three patients, in three comparison settings. Two treatment naive tumors (from initial laparoscopy prior to chemotherapy) were obtained from the peritoneum of patient EOC60 (EOC60-per) and EOC1133 (EOC1133-per), whereas the other three are from interval debulking surgery after three cycles of chemotherapy: one from bowel mesentery of patient EOC60 (EOC60i-meso), and the others from the omentum and ovary of patient EOC868 (EOC868iome and EOC868i-ov, respectively). We compared the methylation between EOC60i-meso and EOC60-per (same patient, treatment naive versus interval, different anatomical site), EOC60-per and EOC1133-per (different patient, treatment naive, same anatomical site), and EOC868i-ome and EOC868i-ov (same patient, interval, different anatomical site). A blood sample from patient EOC868 (EOC868-WBC) was used as a normal control where applicable.

The DNA methylomes were profiled using Agilent SureSelect^XT^ Human Methyl-Seq kit (Agilent Technologies, CA, USA) covering 3.7 M CpGs in 84 Mb target followed by paired-end sequencing with Illumina HiSeq 2500 (Illumina Inc., CA, USA). After sequencing, the methylation patterns at the spanned CpG sites were used for further analysis. There were about 8.64 M such sites, and about 3.07 M of the sites where both compared samples featured coverage of 5x or more. The 5× coverage filtering was performed to prune out low-quality regions, as controlling the number of false positives is sensitive to these. The average per-site read coverage before (after) the filtering was about 11.4× (30.4×). Further details are given in Supplementary material.

#### 3.2.2 Comparing differential methylation calls

The differential methylation calls within each sample pair were obtained as follows. The acquired p-values for sitewise comparisons were adjusted for false discovery rate using the Benjamini-Hochberg procedure (Benjamini and Hochberg, 1995) and calls were made at significance level 0.05, following Lister *et al.* (2009). We used Fisher’s exact test (Fisher) and MOABS (Sun *et al*., 2014), which cannot use a control sample; mixture-adjusted Fisher’s exact test (mix-F; see Supplementary material), InfiniumPurify (Zheng *et al*., 2017), and DSS (Feng *et al*., 2014), which require a control sample; and our method (ML) with a window size of *w* = 1, which operates either with or without a control sample. Further, the mixture-adjusted Fisher’s test and DSS require external tumor purity estimates, which we derived from the corresponding whole-genome sequencing data (WGS) using ASCAT (Van Loo *et al*., 2010).

The heatmaps in Figure 3 display the number of mismatches of either type between any two methods. In more than 91% of the calls, any two methods agree. The results indicate that in all cases, the Fisher’s exact test based methods share most common calls between themselves rather than with the other methods and vice versa, suggesting that the presence or absence of control is not as important as the choice of the method. Typically, a variant lacking a control makes fewer calls than a variant with one, which is expected as the former lacks statistical power as shown in our simulations. A problem characteristic to our samples is that Fisher, MOABS, mix-F lack confidence to make calls between the samples extracted from a single patient, highlighting the importance of tumor purity adjustment. For mix-F and DSS, an appropriate control enables to avoid this problem, which is available in the EOC868i-ome versus EOC868i-ov comparison. Meanwhile, as InfiniumPurify was designed for microarray data, it does not model the heteroscedasticity resulting from read depth differences, so the calls likely correlate with large effect sizes rather than with overall statistical evidence and thus occur at different sites, which would explain why it differs from all the other methods in most cases. In all comparisons, the ML methods produce novel putative differentially methylated sites regardless whether a control is provided or not. In some instances, putative false positives called by especially the methods lacking tumor purity adjustment are pruned. Our hypothesis is that the tumor purity is so low that the accuracy of the Fisher-based methods starts to deteriorate significantly (cf. Figure 2). Finally, in the EOC868i-ome versus EOC868i-ov comparison, the ML based methods result in similar results, suggesting that in the case lacking a control, the methylome of the non-cancer cells is accurately estimated. As the other comparisons do not exhibit such property, the blood sample of patient EOC868 is likely an inaccurate representative of the normal cell methylome for these comparisons and the control-free comparison should be preferred. Findings in known ovarian cancer related genes are summarized in Supplementary material.

**Figure 3:**
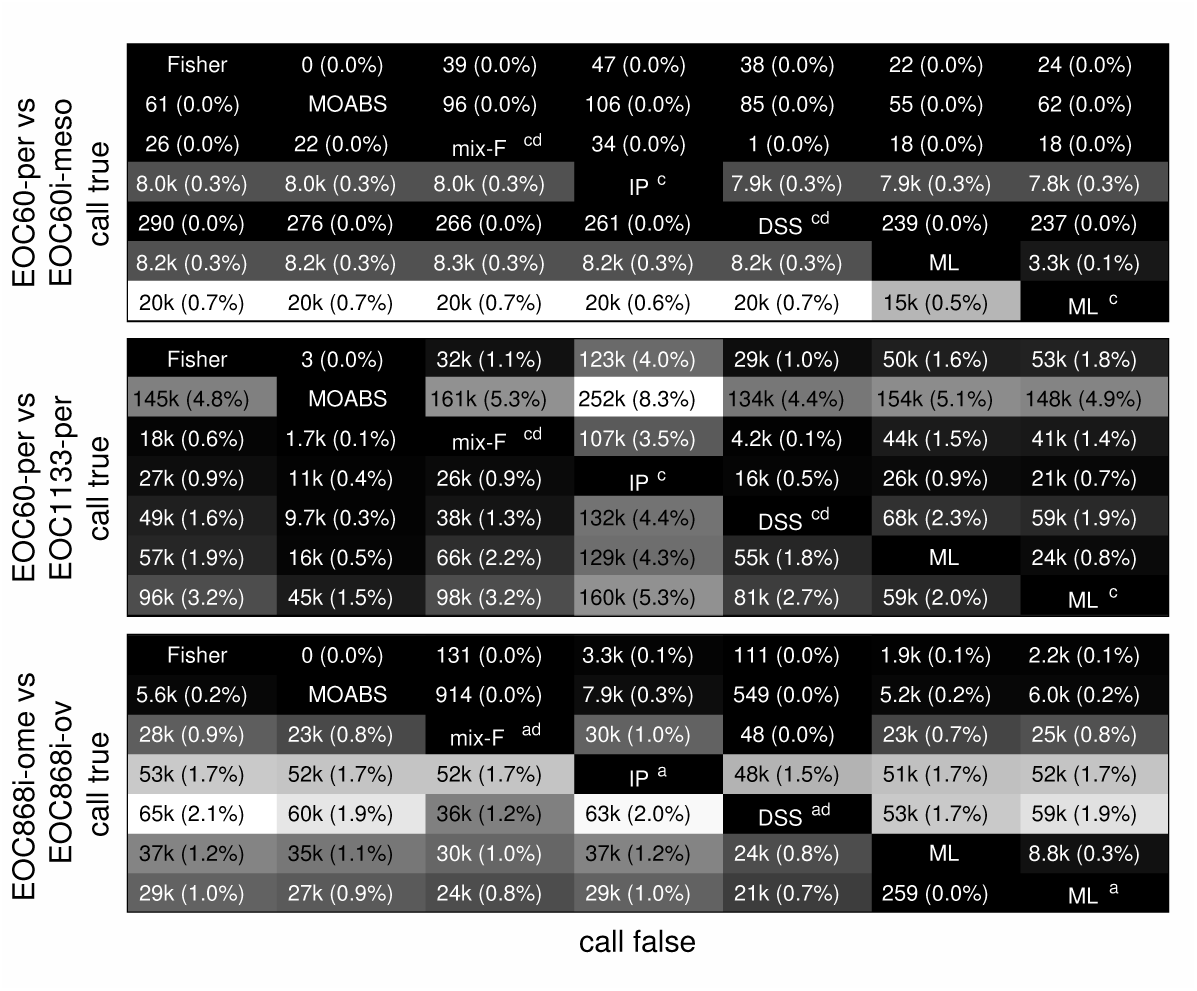
Number/fraction of differential methylation calls in the ovarian cancer samples where the methods differ. Methods: Fisher’s exact test (Fisher), MOABS, mixture-adjusted Fisher’s exact test (mix-F), InfiniumPurify (IP), DSS, and our maximum likelihood method with window size of *w* = 1. Methods with a use a matched control sample, ^c^ use an unmatched control, and d use ASCAT purity estimates from WGS data. The upper triangle shows number of false positives, while the lower triangle shows true negatives.

The computational resources used for the analysis are summarized in Table S1, which indicate that our ML method has a competitive runtime and memory usage when compared to the alternatives. We note that our method has runtime and memory usage that scales linearly with number of CpG sites (see Models and methods), suggesting that even much larger analyses can be done using modest resources.

#### 3.2.3 Comparing purity estimates

To validate the accuracy of our purity estimates, we compared our method with ASCAT (Van Loo *et al*., 2010) and ABSOLUTE (Carter *et al*., 2012), which estimate the tumor purities from whole-genome sequencing (WGS) data. In addition, we used InfiniumPurify to estimate the purities from the same methylation data that was analyzed by our method. The purities estimated using ASCAT and ABSOLUTE from the WGS data and InfiniumPurify from the TBS data of our HGSOC samples are shown in Table S2 with the mean and standard deviation in each comparison. The numbers suggest that the samples vary both in tumor purity and in the degree that the purities differ between the two samples, which explains the varying degree of agreement in the differential methylation calls between our ML method and the alternatives.

Next, we compared the above tumor purity estimates to those obtained using our method. The estimates obtained from the methylation data using our method are shown in Table S2, and they tend to follow the other estimates. To test the reliability of our estimates and if they agree with those reported by the other methods, we estimated the parameters from random substrings of the data. For this, 1,000 uniform random 1,000-site regions of consecutive CpG sites were selected for independent parameter estimation. Figure 4 shows the distribution of estimated parameters for the random substrings of each tumor sample using the methylation data and our method. To test the agreement between the estimates produced by ASCAT, ABSOLUTE, or InfiniumPurify and the estimate by our method, we computed the p-value of obtaining each estimate under the null hypothesis that they follow the distribution of distances from our estimate specified by the random substring estimates, as shown in Table S2. To conclude, we found no evidence that the estimates produced by our method are in disagreement with any of the ASCAT or InfiniumPurify estimates. However, we found some evidence that the ABSOLUTE estimates for the EOC868i-ome and EOC868i-ov samples might differ from those produced by the other methods (p-values ~0.03 when compared with the ML method), which might suggest ABSOLUTE performs better than ASCAT, InfiniumPurify, and our method in low (< 20%) tumor purity settings. In addition, the results indicate that the estimated error parameter is small (less than 0.10 in 77% of the cases) in each case (with no constraints), suggesting that the mixture model explains majority of the data well.

**Figure 4:**
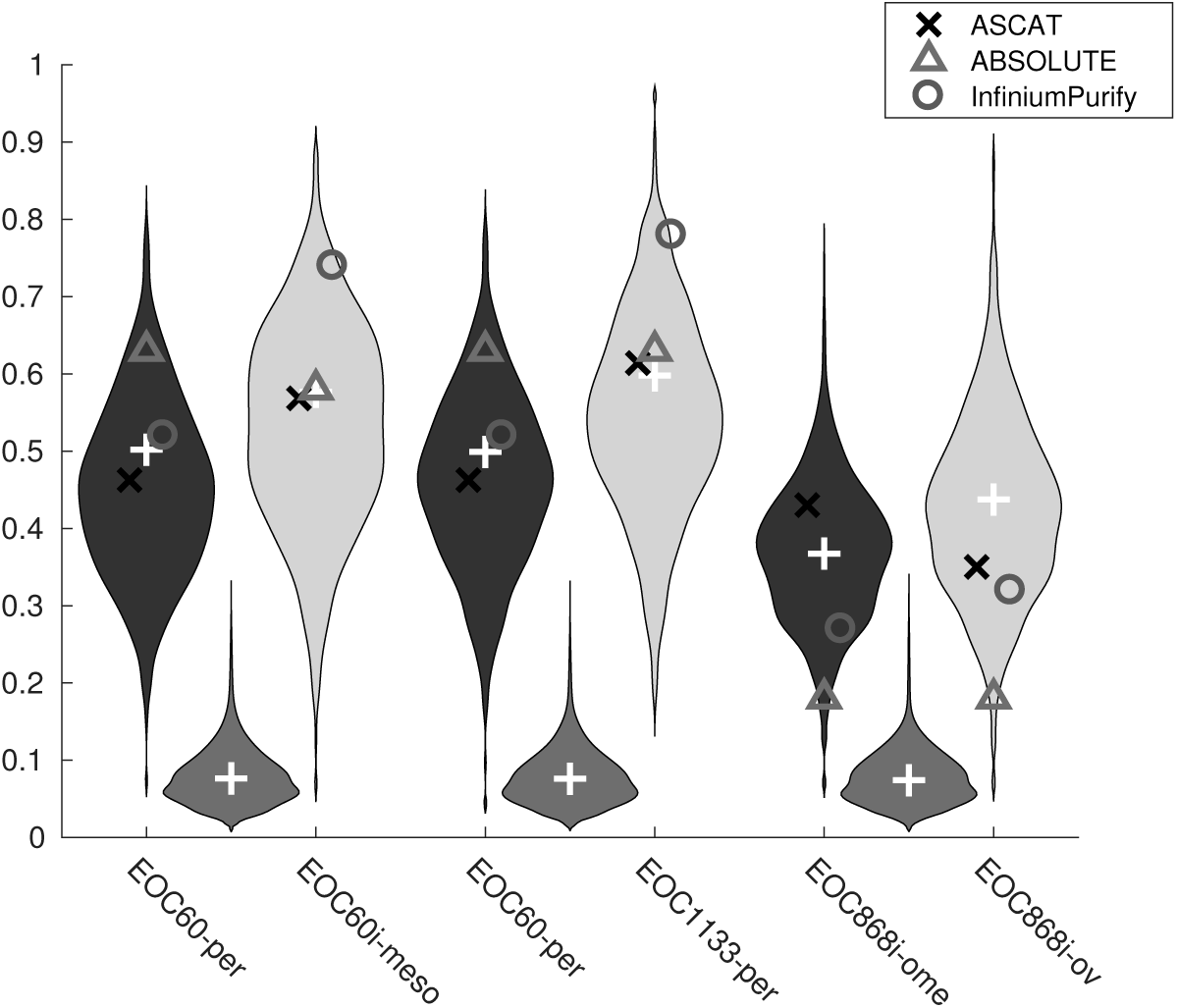
Parameter estimates and an estimate of their variability. Parameter estimates from 1, 000 pieces of 1,000 neighboring sites (violins), the estimates from the whole genome (white crosses), and the tumor purity estimates from WGS data using ASCAT, ABSOLUTE, and from methylation data using InfiniumPurify. Darker and lighter violins correspond to the purities of the compared samples, and the intermediate the (common) error rate.

#### 3.2.4 Differential methylation as a predictor of differential gene expression

To test whether the differential methylation calls produced by our method are more accurate than those of the reference methods, we tested whether our method better predicts gene expression data. For more accurate DNA methylome quantification, we expect to see a stronger correlation between the predicted differential methylation in promoter regions and the corresponding difference in gene expression.

For this analysis, we quantified the Spearman’s rank correlation between the RNA expression ratio and the estimate difference in the DNA methylation in the compared samples. For each method, the difference in the average methylation corrected using the ASCAT purity estimate and the average methylation of the control were used (i.e. solving *y* = (1 – *α*_*j*_) *y*_0_ + *α*_*j*_ *m* for *m* where *y*_0_, *y* are the observed average methylation of the control and the tumor sample, respectively, *α* is the purity, and *m* is the “purified” average methylation), setting the prediction to zero where no call was made. Similarly, the expression values were corrected using the ASCAT purity estimate and a control pooled from all the samples (median). The usage of rank correlation ensures that batch effects and nonlinear relationships between the covariates do not affect the results. The expression data were obtained by analyzing the corresponding RNA-seq data extracted from the tumor samples.

The correlation was quantified for the calls found in the [–1500, +500) region (i.e. the promoter and the first exon area) about the transcription start sites (TSS) of known genes (annotations extracted Jan 2017 from Ensembl (Yates *et al*., 2016)). Hypermethylation in these areas often results in suppressing expression (Witte *et al*., 2014). Outside of this region, no significant correlation was detected in a genome-wide scale. Coincidentally, the effects of global hypomethylation and more focal alterations are expected to be less visible in a genome-wide correlation.

To compare two methods, we quantified the correlation for CpG sites called by either of the two methods. A t-test was used to determine the presence of significant correlation (i.e. if the correlations are nonzero), and Fisher’s transform and a z-test to determine if there is a significant difference between two correlation values (i.e. if two potentially non-zero correlation values differ significantly). The results comparing Fisher’s exact test, MOABS, InfiniumPurify, and DSS to our ML method with a control are shown in Table 1. In each comparison, our ML predictor results in a significant anticorrelation (~–8 to –12%) between differential promoter methylation and the ratio of the expression levels in each sample pair. The results for DSS suggest a similar pattern. For others, significant correlation only exist in some samples. More importantly, our ML predictor results in a stronger correlation, which was found to be significant except against DSS (at significance level 0.05), suggesting that our method results in a more accurate recovery of the methylome.

**Table 1:** Rank correlation between the predicted differential methylation and the differential expression ratio The blocks contain comparison between the reference method and our maximum-likelihood (ML) method. The correlations were computed from the sites where one (or both) of the compared methods make a call. The table lists the correlation for the reference method (Corr ‐) and the ML method (Corr ML), p-values for the hypotheses that these correlations are zero (P_-,0_, P_ML,0_), and p-values for the hypotheses that these correlations are equal (P_-,ML_).

| Sample A/B | Method | Corr - | $P_{-,0}$ | Corr ML | $P_{ML,0}$ | $P_{-,ML}$ |
| --- | --- | --- | --- | --- | --- | --- |
| 60-per /60i-meso | F | -3.6% | 0.04 | -8.6% | $8.4 \times 10^{-7}$ | 0.04 |
| 60-per /1133-per | F | -6.6% | $9.7 \times 10^{-27}$ | -10.5% | $1.5 \times 10^{-64}$ | $9.2 \times 10^{-6}$ |
| 868i-ome/868i-ov | F | -1.9% | 0.21 | -12.0% | $4.7 \times 10^{-16}$ | $1.2 \times 10^{-6}$ |
| 60-per /60i-meso | MOABS | -2.6% | 0.14 | -8.5% | $8.5 \times 10^{-7}$ | 0.02 |
| 60-per /1133-per | MOABS | -6.8% | $1.6 \times 10^{-38}$ | -8.9% | $8.1 \times 10^{-65}$ | $4.4 \times 10^{-3}$ |
| 868i-ome/868i-ov | MOABS | -0.6% | 0.69 | -11.5% | $4.3 \times 10^{-16}$ | $4.4 \times 10^{-8}$ |
| 60-per /60i-meso | IP | -3.2% | 0.04 | -7.7% | $7.1 \times 10^{-7}$ | 0.04 |
| 60-per /1133-per | IP | -5.8% | $1.7 \times 10^{-17}$ | -11.5% | $4.4 \times 10^{-65}$ | $1.6 \times 10^{-9}$ |
| 868i-ome/868i-ov | IP | -3.5% | $5.3 \times 10^{-4}$ | -8.4% | $1.5 \times 10^{-16}$ | $7.0 \times 10^{-4}$ |
| 60-per /60i-meso | DSS | -4.6% | $7.5 \times 10^{-3}$ | -8.5% | $8.4 \times 10^{-7}$ | 0.11 |
| 60-per /1133-per | DSS | -10.3% | $1.7 \times 10^{-63}$ | -10.4% | $3.4 \times 10^{-64}$ | 0.95 |
| 868i-ome/868i-ov | DSS | -9.3% | $9.8 \times 10^{-26}$ | -7.3% | $1.9 \times 10^{-16}$ | 0.11 |

The correlation as a function of the distance from the TSS for the two methods in each of the three cases are shown in Figure S4. The ML method indicates overall stronger correlation than most of the alternatives in the promoter region, which is in agreement with Table 1. The regions outside of [–1500, +500) tend to be poorly correlated with the expression data in a genome-wide level and/or lack data to sufficiently detect one. Our method reveals that changes in the promoter methylation in regions in the vicinity of the TSS and a region at about —800 are strongly reflected in the differential expression of the genes in a genomewide level.

### 3.3 Lymphoma dataset

In addition to the ovarian cancer data, we applied the methods on samples collected from diffuse large B-cell lymphoma (DLBCL) patients. DLBCL is a cancer of B-cells, a type of white blood cell responsible for antibody production, and most commonly derives from mature B-cells (Kuppers *et al*., 1999). DLBCL is the most common type of non-Hodgkin lymphoma in adults, with ~ 7 reported cases per 100,000 people annually.

We used these data in addition to the ovarian cancer data for the following reasons. First, DLBCL is a different type of cancer, so the prevalence and patterns of methylation aberrations likely differs (Witte *et al*., 2014). Second, the methylomes of these samples were profiled using reduced representation bisulfite sequencing (RRBS), which is a different targeting method than used for our ovarian cancer samples. Third, these samples feature a widely different tumor cell fraction (about 84.5% on average versus 42.2% of the ovarian cancer samples according to ASCAT). The analysis is described in detail in Supplementary material.

Again, the results obtained on these data largely agree between the methods (for more than 83% of the calls, any two methods agree; Figure S7, Figure S9–Figure S11). However, provided that our methods are more accurate — as suggested by our simulations and the analysis of the ovarian cancer data — our methods allow ruling out a sizable fraction of false positive calls, and, when used with a control sample, suggest novel findings which the previous methods were unable to identify. Interestingly, with the DLBCL samples the nature of the control sample does not seem to drastically affect identifying the false positives, so even an unmatched control derived from a different tissue, other patients, or from a public database is suitable for these data to gain some of the improvements. The purity estimates produced by our method are consistent with the ones produced by ASCAT and ABSOLUTE from WGS data, or InfiniumPurify from the methylation data (Figure S8).

## 4 Conclusion

We introduced a method that estimates differential methylation between multiple cancer samples featuring varying, unknown tumor purity. The method does not require a prior estimate of the sample purities, nor the methylome of a normal sample, but can estimate these in the process. However, if these information are available, they can be communicated to the estimator in order to improve its accuracy.

The developed method is expected to be of paramount value toward personalized medicine applications, where prognosis, treatment decisions, and response follow-up need to be done using multiple samples, harvested from multiple locations and time points at the individual patient level. Such approach is fundamental for studying heterogeneous diseases, where the comparison can only be done at the patient or small, stratified subgroup level. Our method is the first that allows comparison between tumor and healthy tissue or blood samples, between primary and metastatic tumor samples, or between samples of different patients under the condition where the sample purities vary and a reliable control is absent.

We used Monte Carlo simulations to demonstrate that the method can operate for a wide range of purities, unlike Fisher’s exact test, MOABS, or other methods which do not model the sample composition. Fisher’s exact test and few other methods like DSS can be adjusted for tumor purities but, unlike our method, the adjustment requires prior knowledge of the purities and a reliable control sample. Our method can also exploit such information, but also of the co-methylation of closely located sites, which is a recognized phenomenon (Lister *et al*., 2009; Eckhardt *et al*., 2006) but is neglected by a sitewise Fisher’s exact test and most alternatives. In general, regardless of the setting, our method outperforms all the compared alternatives for a wide range of sensitivity versus specificity.

We applied the method on targeted bisulfite sequencing data from ovarian cancer patients. The results data largely agree between the methods. However, our method suggested up to 5% novel differentially methylated sites, which the previous methods were unable to identify, and allowed ruling out a fraction of false positives. The superior accuracy of our method was confirmed by predicting RNA expression data analyzed with independent methods, and the tumor purity estimates were validated using independent methods using both whole-genome sequencing data (ASCAT and ABSOLUTE) and the methylation data (InfiniumPurify). Thus, we expect that the performance of our method for tumor purity estimation is comparable to the WGS-based methods as well.

The methods were also applied on DLBCL patient data, demonstrating that our methods adapt well to different cancer types and a wide range of tumor purities, and that our method can work robustly with controls derived from various sources. The analyses also demonstrate that our methods can be deployed in genome scale (up to ~ 450 M samples per comparison) and work robustly both with and without a normal cell control.

As profiling DNA methylation in a genome-wide scale has been enabled only recently, computational methods are being developed in order to analyze these data in a meaningful sense. Our method is a significant contribution to this effort for several reasons: First, as the method can estimate all model parameters simultaneously, it requires no additional measurements to supply any configuration parameters. Second, the method can operate either with or without a control sample, which is critical as a control is often unavailable, unreliable, or of suspect. Third, more accurate methods are necessary to elucidate less prominent differences; alternatively, fewer data are needed for equivalent power. Due to these and the important role of DNA methylation, we expect our method to greatly benefit research on cancer and other complex diseases.

The proposed methodology generalizes for estimating differences in any type of sequences, such as mutations in unconverted DNA or DNA copy-number variations. Meanwhile, our methods readily support comparison of multiple samples and more complex sample compositions than used here, provided that the problem is identifiable. Taken together, we believe that our methods enjoy wide applicability in analyzing and comparing measurement data from various sequencing-based platforms.

## References

1. Altman, A. D., Nelson, G. S., Ghatage, P., et al. (2013). The diagnostic utility of TP53 and CDKN2A to distinguish ovarian high-grade serous carcinoma from low-grade serous ovarian tumors. Mod. Pathol., 26(9), 1255–1263.

2. Aran, D., Sirota, M., and Butte, A. J. (2015). Systematic pancancer analysis of tumour purity. Nat. Commun., 6, 8971.

3. Assenov, Y., Muller, F., Lutsik, P., et al. (2016). Comprehensive analysis of DNA methylation data with RnBeads. Nat. Methods, 11(11), 1138–1140.

4. Benjamini, Y. and Hochberg, Y. (1995). Controlling the false discovery rate: A practical and powerful approach to multiple testing. J. Royal Stat. Soc. B, 57(1), 289–300.

5. Berns, E. M. J. J. and Bowtell, D. D. (2012). The changing view of high-grade serous ovarian cancer. Cancer Res., 72(11), 2701–2704.

6. Carter, S. L., Cibulskis, K., Helman, E., et al. (2012). Absolute quantification of somatic DNA alterations in human cancer. Nat. Biotechnol., 30(5), 413–421.

7. Ciriello, G., Miller, M. L., Aksoy, B. A., et al. (2013). Emerging landscape of oncogenic signatures across human cancers. Nat. Genet., 45(10), 1127–1133.

8. Eckhardt, F., Lewin, J., Cortese, R., et al. (2006). DNA methylation profiling of human chromosomes 6, 20 and 22. Nat. Genet., 38(12), 1378–1385.

9. Feng, H., Conneely, K. N., and Wu, H. (2014). A Bayesian hierarchical model to detect differentially methylated loci from single nucleotide resolution sequencing data. Nucl. Acids Res., 42(8), e69.

10. Frommer, M., McDonald, L. E., Millar, D. S., et al. (1992). A genomic sequencing protocol that yields a positive display of 5-methylcytosine residues in individual DNA strands. Proc. Natl. Acad. Sci. U.S.A., 89(5), 1827–1831.

11. Hanahan, D. and Weinberg, R. A. (2011). Hallmarks of cancer: the next generation. Cell, 144(5), 646–674.

12. Hansen, K. D., Langmead, B., and Irizarry, R. A. (2012). BSmooth: from whole genome bisulfite sequencing reads to differentially methylated regions. Genome Biol., 13(10), R83.

13. Harris, R. A., Wang, T., Coarfa, C., et al. (2010). Comparison of sequencing-based methods to profile DNA methylation and identification of monoallelic epigenetic modifications. Nat. Biotechnol., 28(10), 1097–1105.

14. Hebestreit, K., Dugas, M., and Klein, H.-U. (2013). Detection of significantly differentially methylated regions in targeted bisulfite sequencing data. Bioinformatics, 29(13), 1647–1653.

15. Houseman, E. A., Accomando, W. P., Koestler, D. C., et al. (2012). DNA methylation arrays as surrogate measures of cell mixture distribution. BMC Bioinf., 13(1), 86.

16. Illingworth, R. S. Bird, A. P. (2009). CpG islands - 'a rough guide’. FEBS Lett., 583(11), 1713–1720.

17. Kuppers, R., Klein, U., Hansmann, M.-L., Rajewsky, K. (1999). Cellular origin of human B-cell lymphomas. N. Engl. J. Med., 341(20), 1520–1529.

18. Lee, E.-J., Luo, J., Wilson, J. M., Shi, H. (2013). Analyzing the cancer methylome through targeted bisulfite sequencing. Cancer Lett., 340(2), 171–178.

19. Lister, R., Pelizzola, M., Dowen, R. H., et al. (2009). Human DNA methylomes at base resolution show widespread epigenomic differences. Nature, 462(7271), 315–322.

20. Oesper, L., Mahmoody, A., Raphael, B. J. (2013). THetA: inferring intra-tumor heterogeneity from high-throughput DNA sequencing data. Genome Biol., 14, R80.

21. Pan, H., Jiang, Y., Boi, M., et al. (2015). Epigenomic evolution in diffuse large B-cell lymphomas. Nat. Commun., 6, 6921.

22. Shen, H. Laird, P. W. (2013). Interplay between the cancer genome and epigenome. Cell, 153(1), 38–55.

23. Sun, D., Xi, Y., Rodriguez, B., et al. (2014). MOABS: Model based analysis of bisulfite sequencing data. Genome Biol., 15, R38.

24. Sun, S. Yu, X. (2016). HMM-Fisher: identifying differential methylation using a hidden Markov model and Fisher’s exact test. Stat. Appl. Genet. Mol. Biol., 15(1), 55–67.

25. The ENCODE Project Consortium (2012). An integrated encyclopedia of DNA elements in the human genome. Nature, 489(7414), 57–74.

26. Timp, W. Feinberg, A. P. (2013). Cancer as a dysregulated epigenome allowing cellular growth advantage at the expense of the host. Nat. Rev. Cancer, 13(7), 497–510.

27. Van Loo, P., Nordgard, S. H., Lingjaerde, O. C., et al. (2010). Allele-specific copy number analysis of tumors. Proc. Natl. Acad. Sci. U.S.A., 107(39), 16910–16915.

28. Wang, X., Gu, J., Hilakivi-Clarke, L., et al. (2016). Dm-bld: differential methylation detection using a hierarchical bayesian model exploiting local dependency. Bioinformatics, 33(2), 161–168.

29. Wei, S. H., Balch, C., Paik, H. H., et al. (2006). Prognostic DNA methylation biomarkers in ovarian cancer. Clin. Cancer Res., 12(9), 2788–2794.

30. Witte, T., Plass, C., Gerhauser, C. (2014). Pan-cancer patterns of DNA methylation. Genome Med., 6(8), 66.

31. Yang, Z., Jones, A., Widschwendter, M., Teschendorff, A. E. (2015). An integrative pan-cancer-wide analysis of epigenetic enzymes reveals universal patterns of epigenomic deregulation in cancer. Genome Biol., 16, 140.

32. Yates, A., Akanni, W., Amode, M. R., et al. (2016). Ensembl 2016. Nucl. Acids Res., 44(D1), D710–D716.

33. Yoshihara, K., Shahmoradgoli, M., Martinez, E., et al. (2013). Inferring tumour purity and stromal and immune cell admixture from expression data. Nat. Commun., 4, 2612.

34. Zheng, X., Zhao, Q., Wu, H.-J., et al. (2014). MethylPurify: tumor purity deconvolution and differential methylation detection from single tumor DNA methylomes. Genome Biol., 15(8), 419.

35. Zheng, X., Zhang, N., Wu, H.-J., Wu, H. (2017). Estimating and accounting for tumor purity in the analysis of dna methylation data from cancer studies. Genome Biol., 18(1), 17.

